# Identification of outcome-related driver mutations in cancer using conditional co-occurrence distributions

**DOI:** 10.1101/075408

**Authors:** Victor Treviño, Emmanuel Martínez-Ledesma, José Tamez-Peña

## Abstract

The methods proposed for the detection of cancer driver mutations are based on the estimation of background mutation rate, impact on protein function, or network influence. Instead, we focus on those influencing patient survival. For this, an approximation of the log-rank test has been systematically applied even though it assumes a large and similar number of patients in both risk groups, which is violated in cancer genomics. Here, we propose VALORATE, a novel algorithm for the estimation of the null distribution for the log-rank test independently of the number of mutations. VALORATE is based on conditional distributions of the co-occurrences between events and mutations. The results using simulations, comparisons with other methods, TCGA and ICGC cancer datasets, and validations, suggests that VALORATE is accurate, fast, and can identify known and novel gene mutations. Our proposal and results may have important implications in cancer biology, in bioinformatics analyses, and ultimately in precision medicine.

## Introduction

Cancer is a genetic disease characterized by the progressive accumulation of mutations ^1^. The recent sequencing technologies are revolutionizing cancer medicine with the rich characterization of genetic mutations ^2^. International efforts, such as The Cancer Genome Atlas (TCGA) and the International Cancer Genome Consortium (ICGC), have been established to scrutinize several cancer types by generating large amounts of cancer genomics data ^3^. Nevertheless, because of its intrinsic complexity, there is a need for more advanced and precise methods for the analysis of these data to gain deeper understand this deadly disease.

One fundamental problem in cancer genomics is the detection of functional mutations. In this context, the progressive accrual of mutations ^1^ and its heterogeneity ^4^ have fueled the theories of clonal expansion ^5^ in which ‘driver’ mutations have functional roles that confer cell fitness advantages whereas ‘passenger’ mutations are the result of the inherent random mutational process ^6,7^. The detection of driver genes is challenging because the observed frequency of gene mutations is relatively low for most of the genes ^8^. Moreover, the detected gene driver mutations explain only a fraction of mutations per patient ^1,8^ suggesting that novel methods to detect driver mutations are still needed even though other genetic alterations may also be present (such as copy number, fusions, and epigenetic alterations).

Most of the methods to detect driver mutations that have been proposed ^9^ can be classified according to its main concept in *(i)* methods that identify recurrent gene mutations, and *(ii)* methods that identify the impact of the disrupted protein function. For recurrent gene mutations, methods such as MutSigCV ^10^ and OncodriveCLUST ^11^ use specific ‘null’ models to estimate background mutation rates and recognize those failing the null model. Indeed, these ideas have been applied also to non-coding regions ^12^. In addition, there is a subclass of methods focused in recurrent gene mutations per network modules such as HotNet2 ^13^. To detect the functional impact, for example, SIFT ^14^ and MutationAssessor ^15^ look at disruptions in evolutionary conserved functional domains, which can be assessed per variant. Contrary, OncodriveFM ^16^ examines a set of variants evaluating whether functional impacts per variant is shifted toward high impacts.

The methods mentioned above consider mutations in a cellular or molecular view where the effects take place locally assumed to improve the fitness of the tumor cell to its microenvironment. Nevertheless, there are gene mutations that violate the model assumptions complicating its identification. For example, when the functional effect of a mutation is unknown, or when a mutation is indeed similar to those evolutionary conserved but that in the human context are damaging ^17^. On the other hand, model parameters may be over-simplified, such as the replication timing that has been shown to be highly influenced in a cell type-specific manner ^18^. In addition, there could be other situations more difficult to model such as mutations appearing in recurrent tumors ^19^, tissue invasion or metastasis ^20^–^22^, or co-occurring alterations ^23^. Some of these will clearly have an effect on tumor development and time to death, metastasis, or recurrence. Therefore, to overcome some of these limitations, here we adopted a ‘long-term’ population risk view in which mutations may influence patient survival. This is important because the precise identification of mutations relevantly associated with clinical outcome could be crucial for treatment and precision medicine ^24^.

Generally, the identification of gene mutations associated with survival is done by forming two risk groups splitting those subjects carrying the mutation from those who not. The differences in time to death between these risk groups is used to detect the relevant mutated genes. The statistical significance of this time difference is usually estimated by computing the Log-Rank statistic followed by the computation of its probability of being zero ^25^. Commonly, this p-value is approximated by Gaussian or χ^2^ distribution ^25^. This procedure, which will be referred as the ‘Approximate Log-Rank Test’ (ALRT), assumes that both populations have a similar number of subjects and that this number is large ^26^. In cancer genomics, however, these assumptions are generally not met because the frequency of mutations in a gene across patients is generally low ^8^. To address these issues, a precise estimation of the null distribution is required. However, this is challenging since there is no analytical form of the log rank statistic (to our knowledge) and the number of combinations is astronomically high even for a low number of subjects and mutations, making exact estimations of the null distribution computationally impractical. Therefore, due of the lack of statistical and computational tools, the ALRT is commonly applied as can be seen in cancer genomics data portals (see *Correlate Clinical vs Mutation* in ‘Clinical Analyses’ from any cancer type within http://firebrowse.org). Recently, a method that accelerates the estimation of the null distribution, ExaLT, has been proposed in which the number of combinations is reduced under certain circumstances controlled by a precision parameter ^27^. However, this algorithm is still prohibitively slow even for moderated population sizes (around n=200) and precision values. Consequently, there is yet a need for fast and accurate methods to estimate the null distribution and the probability of associations of between mutated genes and cancer survival time, which may help the discovery of novel genes, mutations, and provide important insights to cancer biology.

Here we propose VALORATE (*Velocity and Accuracy for the LOg RAnk TEst*), a novel algorithm, which quickly provides a precise and accurate estimation of the empirical null distribution and the probability value of the Log-Rank statistic being zero regardless of the population size and the fraction of mutations. We first validate the accuracy and velocity of VALORATE in simulated and cancer data. Then, we apply VALORATE to analyze the gene mutations associated with survival times in 61 cancer datasets including more than 40 cancer types from TCGA and ICGC, which cover 11,655 and 2,779 cancer samples respectively. We note that, regardless of the method, the significance can be influenced by hypermutated samples. Next, we show comparisons of the estimations of VALORATE and the ALRT suggesting that the ALRT may detect many false positives and false negatives genes. As a consequence of these differences, we observe that the proportions of genes associated with low- and high-risk groups are largely different. We find that genes associated with survival using VALORATE were mostly cancer-type specific. From the identified genes, many are well-known cancer genes but many others are novel associations. These results seem to be reliable because significant genes appear to be expressed in the mutated tissues and its mutations have high functional impacts. We conclude that VALORATE is a valuable tool for cancer genomics and may be useful for other statistical applications.

## Results

### Validation of the VALORATE algorithm

The VALORATE algorithm shown in Figure 1 (see details in Materials and Methods) is based on the postulate that the log-rank distribution *L* will be highly dependent on *k*, the number of co-occurrences of events and mutations when the number of subjects in one group is very low or presumably when there is a highly unbalanced number of subjects between groups. Thus, *L* is a weighted sum of conditional distributions *L*_*k*_. The procedure is fast to compute because *L*_*k*_ can be estimated by sampling.

**Figure 1.**
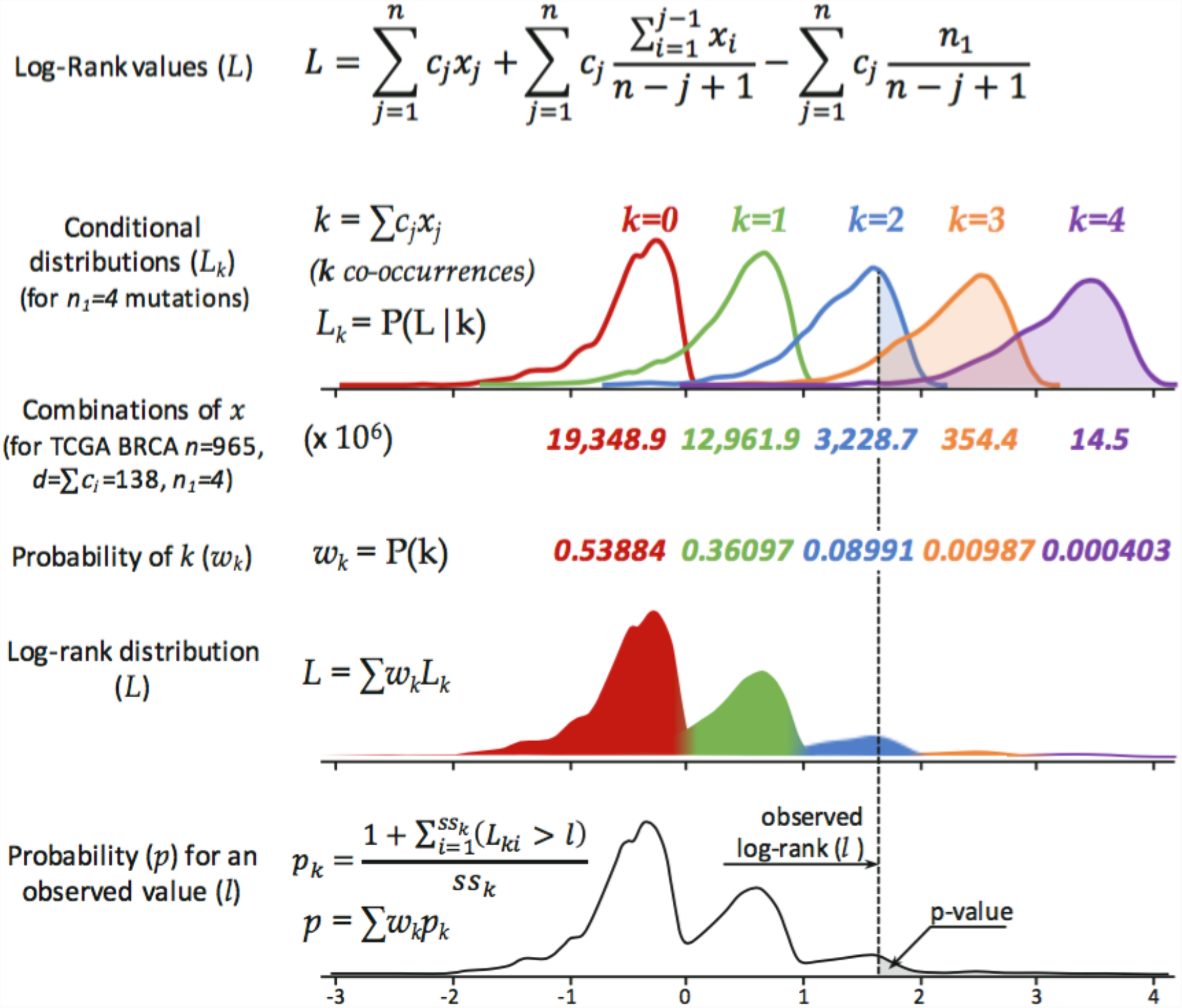
Overview of the VALORATE algorithm. For a dataset having *n* samples, *d* deaths, and a gene having *n*_1_ sample mutations coded in the vector *x* of mutated subjects, the conditional distributions *L*_*k*_ are estimated by random sampling *x* over *k*, where *k* is the number of co-occurrences (events that are also mutated). The proportional weight of each *L*_*k*_ can be estimated by the contribution to the total number of combinations, which for a given *k* can be calculated by *C*(*n-d*, *n*_1_-*k*)\**C(d, k)*, where *C* is the combination function. The overall distribution is then estimated by a weighted sum on *L*_*k*_. Finally, the p-value for an observed log-rank value in a mutated gene can be estimated by weighting the conditional p values over *k*.

To show the accuracy of VALORATE in the estimation of the log-rank distribution, we used simulations comparing the exact distribution of all possible combinations of the log-rank statistic with the distribution estimated by VALORATE. A representative simulation shown in Figure 2A suggests that VALORATE can accurately estimate the log-rank distribution, and it is consistent in a variety of simulated scenarios (Supplementary Figure 1). As expected, the most extreme statistics of the distribution were not observed due to random sub-sampling (Supplementary Figure 2). This small caveat is not an issue because the estimated *p* values according to our procedure for these extreme statistic values would be, correctly, close to 0 and the observed statistics close to these extremes are accurately sampled (Figure 2B, and Supplementary Figure 1B).

**Figure 2.**
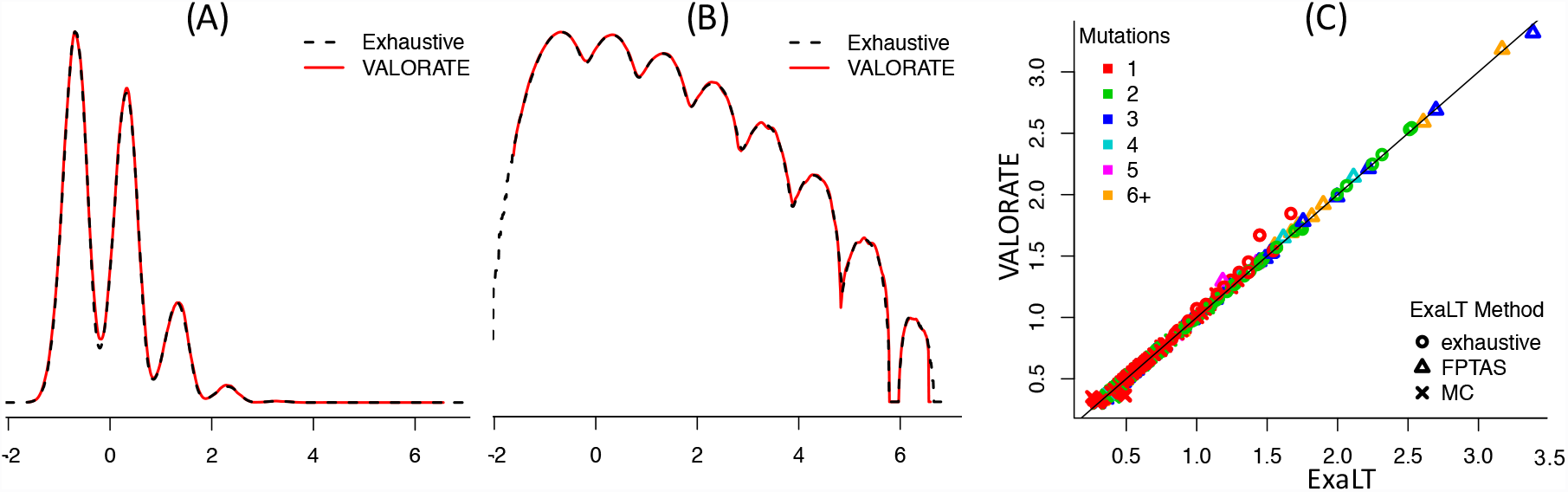
Accuracy of VALORATE. A) Comparison of the exact log-rank distribution for a simulated dataset with the distribution estimated by VALORATE using *ss=100,000*. The simulation was estimated by *n=100* subjects, *d=10* events, and *n_1_=7* mutations, which generates 16,007,560,800 combinations. (B) Densities in logarithm base 10 to show details in low-density regions (above 3). (C) Comparison of the p-values estimated by VALORATE and those estimated by ExaLT for the GBM dataset from Vandin *et al.* (2015) shown in supplementary material (TableR.txt file in https://github.com/fvandin/ExaLT). p-values are shown in negative of the logarithm base 10.

To show the accuracy of the VALORATE procedure regarding the p-value estimations, we ran some simulations. The results show that when the assumption of a similar number of subjects in the two groups are met (*n=100* subjects and *n*_1_=50 mutated), as in the ALRT, the p-values estimated by the ALRT and VALORATE are highly similar and highly correlated independently of the co-occurrence *k* (Supplementary Figure 3). However, when the number of subjects between groups becomes more dissimilar (*n_1_= 30, 14, or 7*), the differences in p-value estimations turn higher, which correlates with changes in the symmetry and shape of the overall log-rank distribution. Moreover, the differences in p-value estimations also depend on the number of events co-occurring in the mutated risk group (Supplementary Figure 3). For example, in an extreme case when *n*_1_=7 where the number of events in the mutated group was *k=0* (so the 7 mutated samples are censored), the ALRT estimated a p-value of 0.15 whereas VALORATE estimated 1.8×10^−4^ (Supplementary Figure 4). Contrary, in a case where *k=1*, the ALRT estimated a p-value of 3.5×10^−6^ whereas VALORATE estimated 0.27. For these estimations, VALORATE used *n*_1_ =7 estimated distributions *L*_*k*_, whereas the ALRT uses one χ^2^ distribution.

**Figure 3.**
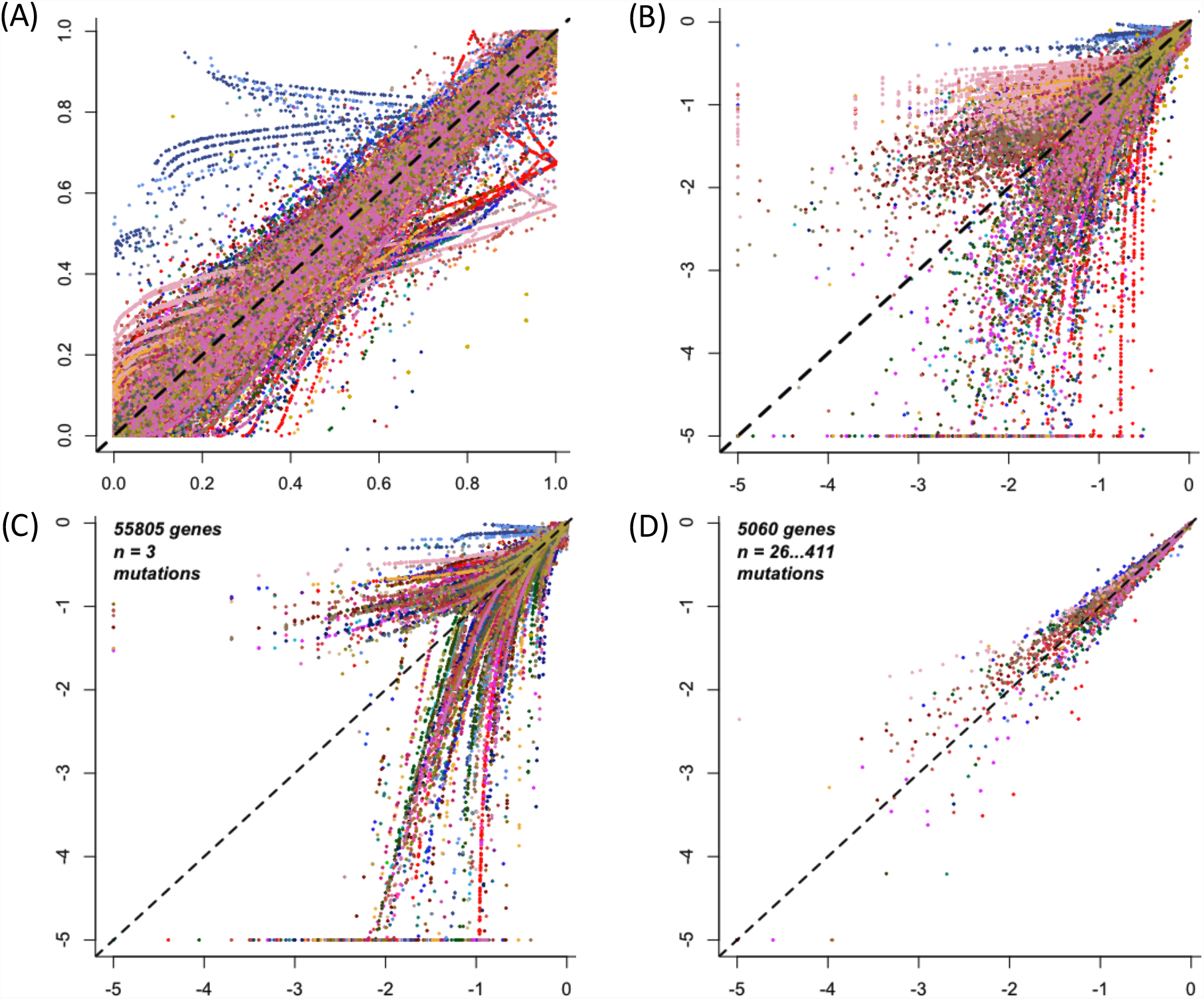
Comparison of the p-value estimations from VALORATE and the ALRT in cancer datasets. (A) p-Value estimated by VALORATE (horizontal axis) and the ALRT (vertical axis). (B) Same than (A) in logarithm base 10 scale to highlight region of significance. Only genes having 4 or more mutations are shown. (C) The p-values for genes having 4 samples mutated. (D) The p-values for genes having more than 25 sample mutated.

The above results indicate that the VALORATE procedure can accurately *(i)* approximate the exact *L* distribution independent of the shape of the log-rank distribution and *(ii)* calculate the correct p-values since they converge to the ALRT when *n*_1_ is similar to *n/2*. This proposes that VALORATE is superior to the ALRT for estimating the probability of the difference between two survival curves, especially in the cases where *n*_1_ departs from *n/2*.

Then, we evaluated the precision of VALORATE on repeated runs across different values of sampling sizes from 10^3^ to 10^6^. The results show that at *ss=10,000*, different runs are almost indistinguishable indicating high precision (Supplementary Figure 5). Even for *ss=1,000*, the shape of the distribution is highly similar to that in *ss=1,000,000* showing that the procedure is consistent and robust. Nevertheless, at low sampling size, two runs may display slight differences. Although the differences in the estimation of the distribution and hence the p-values should be small, it is preferable to use larger sampling sizes to avoid small fluctuations between runs. Therefore, we used *ss=100,000* for the cancer data analyses.

To evaluate VALORATE in cancer data including the estimation of p-values, we compared the calculations against those provided by ExaLT, which is based on three different approaches. The results are highly similar between different methods and number of mutations (Figure 2C) suggesting that the p-value estimations from VALORATE are also accurate in cancer data.

The computation time is an important issue because genomics data is being generated at high rates and typical analyses may involve estimations for the available stratifications (e.g. cancer sub-types, hormonal status, histological grades) and within systematic pipelines and data versions. Thus, we assessed the running time for VALORATE and ExaLT changing the main parameter associated with accuracy. Even that both algorithms are different in essence, this test illustrates the time scale needed and how it grows. The Table 1 shows that VALORATE run more than 10,000 times faster than ExaLT. Besides, the running time of VALORATE does not grow drastically.

**Table 1.** Running time for the probability calculation of 4 genes in two algorithms.

| Accuracy | VALORATE |  |  | ExaLT (FPTAS) |  |  |
| --- | --- | --- | --- | --- | --- | --- |
|  | Parameter<br>(ss) | CPU Time<br>(sec) | Time<br>Growth | Parameter<br>(af*) | CPU Time<br>(sec) | Time<br>Growth |
| **Low** | 1,000 | 0.10 |  | 1,000 | 563 |  |
| **Modest** | 10,000 | 0.15 | 1.5x | 100 | 710 | 1.3x |
| **Good** | 100,000 | 0.37 | 2.5x | 10 | 2,405 | 3.4x |
| **High** | 1,000,000 | 2.24 | 6.0x | 1 | 34,910 | 14.5x |
| **Extreme** | 10,000,000 | 14.28 | 6.4x | - | - |  |

\* Approximation factor

### Comparison of detected gene mutations

Recent studies have suggested that the ALRT may provide different and false results for the identification of cancer gene mutations associated with outcome ^27^. We used VALORATE to compare and analyze the implications in the use of a more appropriate test in 61 cancer datasets from the TCGA and ICGC (Supplementary Table 1) covering 14,434 samples, 528,124 genes and 3,103,054 mutations. We first obtained the p-values using the ALRT compared against those provided by VALORATE for the 152,466 genes having more than 3 samples mutated in any cancer. The results demonstrate that, overall, the p-values are highly correlated (Figure 3A), which further support the estimations provided by VALORATE. Nevertheless, a more specific analysis over the most significant region (p < 0.1) shows that the estimations can be notably different (Figure 3B). Only 148 genes (0.097%) show p-values < 0.001 in both tests. At p < 0.01, the ALRT seems to call 1.8 times more genes as significant compared to VALORATE. However, we observed some differences across cancer datasets (Supplementary Figure 6). For example, some cancer types show more detections in the ALRT such as breast cancer (BRCA) and others cancer types show more detections in VALORATE such as uterine corpus endometrial carcinoma (UCEC). We then observed that the differences in p-value estimations between the ALRT and VALORATE are specific for genes having few number of mutations (Figure 3C-D), where the ALRT assumptions are not met. Indeed, the differences in p-value estimations decrease for increasing number of mutations (Supplementary Figure 7). This indicates that, as the simulations suggested, the use of the ALRT estimation is progressively detrimental when decreasing the number of mutations on cancer data.

### Identification of outcome associated driver mutations in cancer

The results shown above may have important implications in cancer genomics because it raises the possibility that other genes can be identified and that some of the previously identified genes using the ALRT could be suspicious. We, therefore, selected the most significant genes after correction by false discovery rate (FDR < 0.333). We observed large differences in the selected number of genes between VALORATE and the ALRT (Supplementary Figure 8A). Only 7% of the genes was identified in both tests or 34% when considered the rank of top genes (Supplementary Figure 9). Two examples of such discrepancies are shown in Figure 4 for the genes RAB42 in breast cancer and LMTK2 in thyroid cancer. The gene RAB42 is the most significant reported by the TCGA in BRCA (p=1×10^−8^, q=9×10^−6^, http://firebrowse.org, doi:10.7908/C10Z72M8), however, in VALORATE the significance is marginal (p=0.02) and not selected after FDR correction (q=0.44). Contrary, the LMTK2 in THCA, which is significantly mutated by frequency using MutSigCV ^10^ from TCGA but not associated with time to death using the ALRT (http://firebrowse.org, doi:10.7908/C1542N2H), is the most significant mutated gene using VALORATE (p=0.00026, q=0.075). All these results suggest that VALORATE can identify genes that are missed by the ALRT and mark genes whose association with survival can be spurious, which contributes to providing important insights in cancer biology.

**Figure 4.**
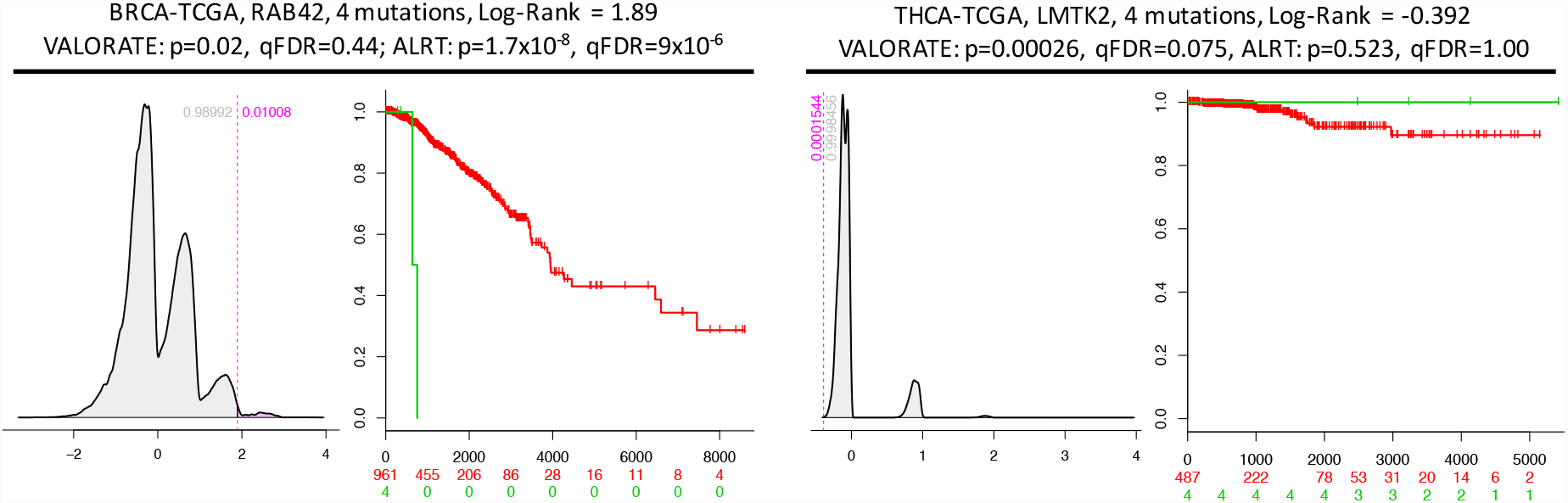
Examples of differences in p-value estimations. (A) RAB42 in BRCA from TCGA.(B) LMTK2 in THCA from TCGA.

Nevertheless, we observed an apparent excess of significant mutations in uterine corpus endometrial carcinoma, bladder, and colon (China) (UCEC, BLCA, COCA-CN respectively) having more than 200 genes associated with survival according to VALORATE (Supplementary Figure 8A). The number of significant genes was weakly associated with dataset characteristics such as the number of samples, events, or censoring (Supplementary Figure 10). Instead, it was clearly associated with hypermutated samples (Supplementary Figure 11), which represent a minor proportion of samples with an exacerbated number of mutations ^28^. This bias could not be observed using the ALRT in UCEC because there were no significant genes after FDR correction. But in breast cancer (BRCA) where the ALRT detects many significant genes and VALORATE does not, the same issue arises using the ALRT (Supplementary Figure 11). This result proposes that hypermutated samples may bias the survival analyses of mutated genes. To explore this further, we re-analyzed all datasets removing the top 5% most mutated samples. The results show a substantial reduction of detected genes (Supplementary Figure 8B) suggesting that indeed hypermutated samples influence the selection of many genes in several cancer types (except gliomas). Therefore, for further analysis, we used a refined criterion to remove hypermutated samples avoiding the removal of samples having few mutated genes (Supplementary Figure 12 and Materials and Methods).

### Significant genes and risk assessment

After removing hypermutated samples and using an FDR=0.333, the ALRT calls 2,445 genes significant while VALORATE calls only 255 (Supplementary Figure 13). This decrease is similar across cancer types. Only in Gliomas, we observed 164 significant genes and the association was not related to hypermutated samples (Supplementary Figure 14). From significant genes, 212 were detected in both tests (Figure 5A). Interestingly, the risk associated with genes between tests is different (Figure 5B, chi-square test=2.2×10^−16^). In the ALRT, only the 2% of identified genes are associated with low risk whereas in VALORATE low-risk genes reach 28%. We observed few but more low-risk gene mutations in cervical squamous cell carcinoma, esophageal carcinoma, glioblastoma, ovarian, stomach-esophagus cancers, thyroid cancer, uterine corpus endometrial cancer and few others (CESC, ESCA, GBM, OV, STES, THCA, and UCEC respectively, see Supplementary Figure 15). These findings may have implications in cancer biology because, apart from few exceptions such as IDH1 in gliomas, most coding mutations detected so far in cancer have been associated with decreasing survival rates using the ALRT ^29^. Moreover, these results may also help to design low-risk biomarkers in other cancer types.

**Figure 5.**
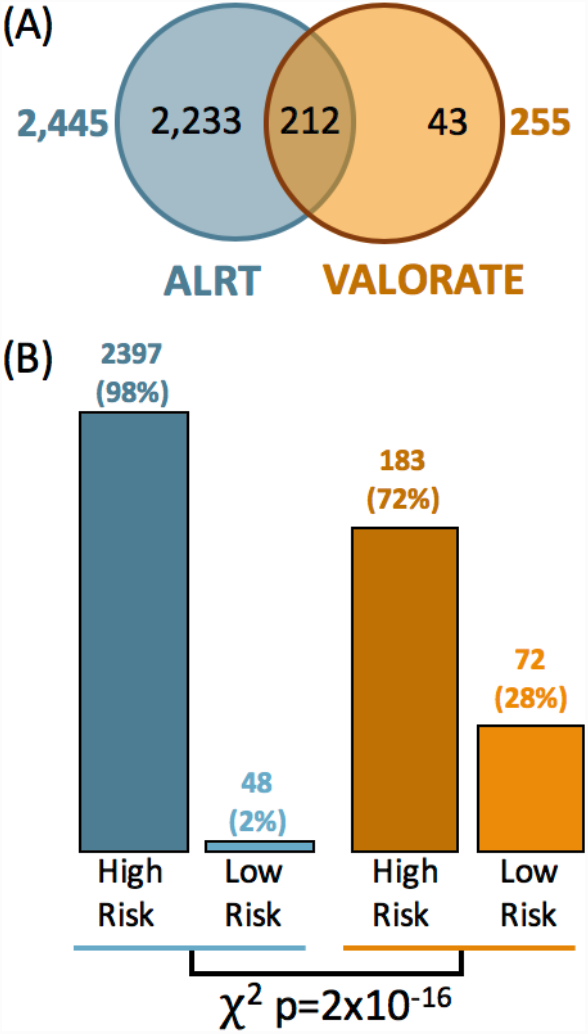
Comparison of significant genes and associated risk. (A) The genes detected in two methods. (B) The number of genes associated with high- and low-risk.

### Significant genes across cancer types

It is expected that significant genes could be associated with survival in several cancer types because it is well known that some genes are broadly mutated such as TP53, PIK3CA, PTEN, KRAS, ARID1A ^8^, and others (see www.tumorportal.org). Thus, we compared the significant genes detected by VALORATE across cancer types (Figure 6). At FDR=0.333, only TP53 and MUC4 were found significantly associated with survival in at least two ‘distinct’ cancer types (TP53 to GBMLGG, LAML, BOCA-FR, BTCA-JP, PRAD-UK, and MUC4 to COCA-CN, KICH, and KIPAN). TP53 was marginally significant (FDR > .33 and p < 0.05) in many other cancer types and MUC4 in a few others. Interestingly, ATRX, well-known in glioblastomas ^30^, was significant in gliomas, neuroblastoma, and pheochromocytoma/paraganglioma, all these related to nervous system. Some genes appear significant in one cancer type and marginally significant in others types. Extreme cases are MUC16, MUC17, PCLO, and DNMT3A that are marginally significant in few more types. Some few genes appear significant in datasets that are composed of similar cancer types such as gliomas (GBMLGG, which is composed of glioblastoma and low-grade gliomas), stomach-esophagus (STES), and kidney (KIPAN). Apart from these few exceptions, the majority of the genes seem cancer-type specific (Figure 6) even when the significance assumption is relaxed (Supplementary Figure 16). Remarkably, we did not observe significant survival association to other well-known cancer genes such as PIK3CA (lowest p=0.007, q=0.5 in UCEC), BRAF (lowest p=0.028, q=0.98 in KIRP), and FBXW7 (lowest p=0.018, q=0.99 in MELA-AU). From the 77 genes in Figure 6, only 12 were also significant in the TCGA systematic analyses using the ALRT. Overall, these results suggest that association to survival is a different feature that mutation frequency supporting the exploration of association to a broader set of mutated genes rather than those detected by MutSigCV or similar methods.

**Figure 6.**
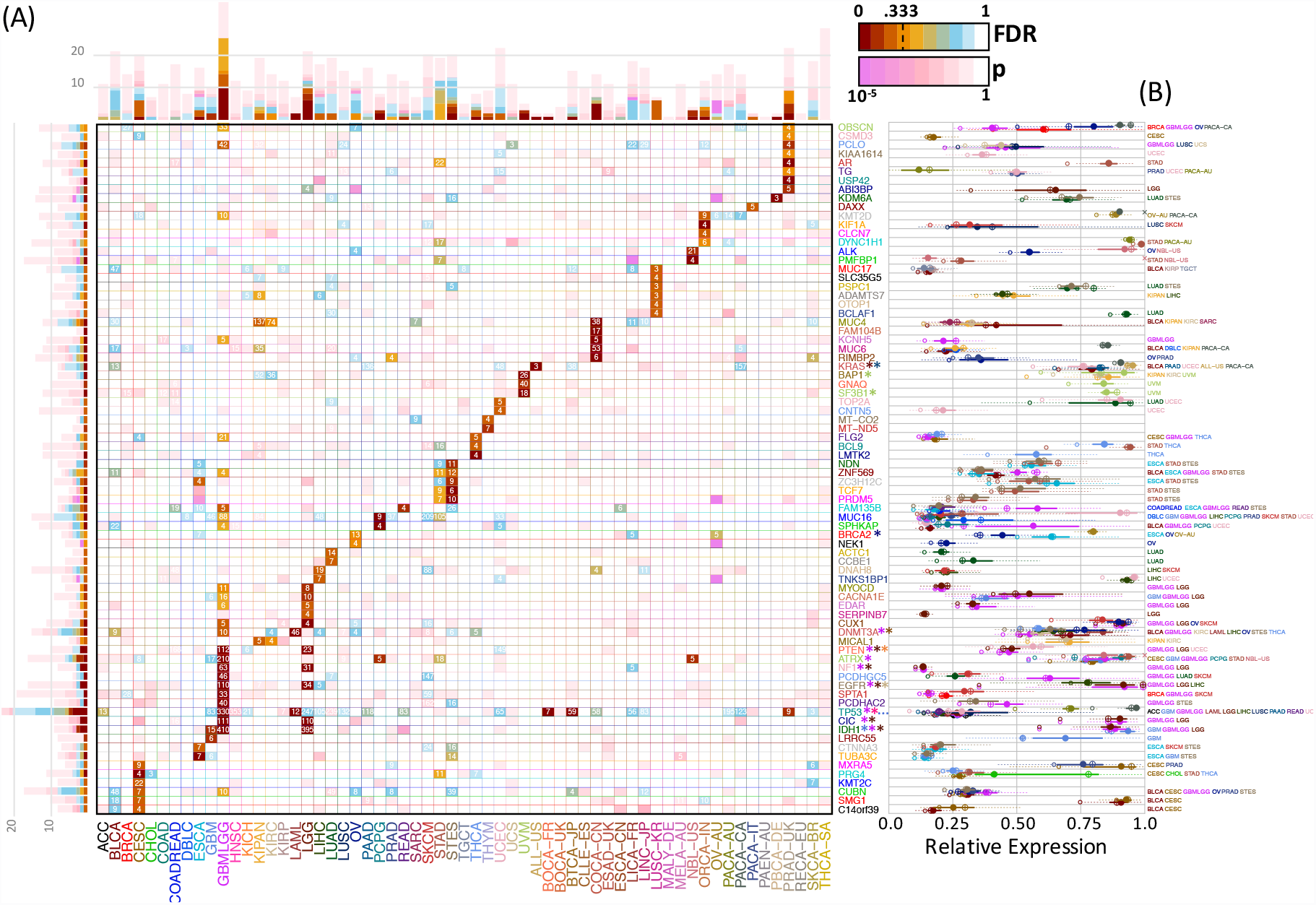
Significant genes across cancer types detected by VALORATE. A) Map of genes detected (vertical) across cancer types (horizontal). The color shades (from brown to orange and cyan) in each cell correspond to FDR q-value when less than 0.9 in which case the number of mutated samples is also shown. In cases where FDR > 0.9, the p-Value is shown in shades of pink. The bars show the accumulated number of cancer types for genes (left bars) or genes for cancer types (top bars). For Gliomas (GBMLGG) and Low-Grade Gliomas (LGG) only the top 10 genes are shown. The genes marked with an asterisk (*) have been reported as significant in FireBrowse (http://firebrowse.org). (B) Gene expression levels of the significant genes in those cancer types whose FDR > 0.9 and whose gene expression values were available. The filled circles represent the median (50%) of the expression level and the continuous line the 25% and 75% using all samples. The crossed circles represent the median (50%) of the expression level and the dotted line the 25% and 75% from the mutated samples. The smaller hollow circles represent the lowest expression value from the mutated samples.

### Functional Validation of Significant Genes

In this study, 43 genes were significant only in VALORATE and can be considered novel associations (Figure 6). In addition, many significant genes were highly ranked in VALORATE and obscured by its rank using the ALRT (Supplementary Figure 9). Therefore, we asked whether the significant genes may have plausible functional roles. For this, we made two tests regarding gene expression and functional impact of mutations. For the first, we reasoned that if a mutation in a coding gene is associated with survival, the gene should be expressed. The Figure 6 shows that most of the significant genes are expressed at some level. Some genes show expression levels around 15% (relative to their cancer) but the expression levels across tissues are highly consistent (for example TP53, MUC17, FLG2, TUBA3C) suggesting a common functional level of expression. Moreover, the expression of mutated samples is highly similar to those observed in its cancer type.

We also tested the possible functional impact associated with the mutations of significant genes that may affect active sites, interactions with other proteins, and 3D folding. This can be corroborated comparing the non-synonymous mutations against conserved sequences in homolog proteins. For this, we used MutationAssesor ^15^, which summarize the functional impact into neutral, low, medium, and high impact. We observed that the mutations in the identified genes have significantly higher impact categories than randomly chosen mutated genes from same cancer types and a similar number of mutations (Supplementary Figure 17).

All these results suggest that, in general, coding mutations of the identified significant genes can be functional.

## Discussion

A fundamental problem in cancer genomics and precision medicine is the determination of genomic alterations that could be associated with survival times. The selected alterations are then the seeds for research studies of biological mechanisms, drug discovery, and possible treatments. The identification of the important genomic alterations is, however, challenging because most of the observed alterations are present in a low number of patients, there are thousands of alterations to test, and the associations need to be tested in several subject strata (grades, hormonal status, molecular subtypes, etc.). It has been recently shown that the statistical approximations used for this identification in cancer genomics are inaccurate ^27^. The failure is basically due to the low number of patients presenting a specific alteration that generates heavily unbalanced population sizes. Although an accurate tool has been recently proposed ^27^, it is prohibitively slow to compute in practice. In this work, we revisit the problem of estimation and propose, VALORATE, a novel estimation procedure that is independent of the number of alterations. We show by simulations that VALORATE is fast, precise, and accurate. In comparison with another method, VALORATE also provided accurate p-values in cancer data. We show that VALORATE is accurate when comparing largely unbalanced populations and highly similar to the ALRT when the populations are balanced. Thus, VALORATE can be used in both cases. This should facilitate its use and implementation in current bioinformatics pipelines (see Code section in Methods).

We demonstrate that the ALRT generates poor results under unbalanced populations. This agrees with previous results^26,27^. Furthermore, our simulations demonstrate that the ALRT overestimates the significance for higher values of co-occurrences (*k*) and underestimates the significance for lower values of co-occurrences even in the same number of mutations (*n*_1_). This could explain the large differences in genes called significant between the ALRT and VALORATE across cancer types.

Using VALORATE, we identified that hypermutated samples may bias the estimation of p-values. This issue is related to two factors. First, VALORATE is a univariate procedure and suffers its same caveats, it tests the alterations in one gene (or locus) at the time being blind to other alterations. Second, the biology of the relation cancer-mutation-patient-survival is complex. In UCEC for example, hypermutated samples show high survival times and censored. Contrary, in BRCA, the hypermutated samples present poor survival. Here, we first removed top 5% most mutated samples to demonstrate the impact of the hypermutated samples. Then, we refined the criteria avoiding the unnecessary removal of samples to yield more fair estimations. Nevertheless, the decision whether to remove samples, how many, and which, deserves attention.

In datasets that are generated by the union of cancer types such as stomach-esophagus (STES), colorectal (COADREAD), pan-kidney (KIPAN) and others, we observed similar p-value estimations of the mutated genes in individual datasets compared to the merged datasets.

However, we observed 164 significant genes in gliomas (GBMLGG) while only 19 and 2 genes were significant in low-grade gliomas (LGG) and glioblastomas (GBM), respectively. This could be related to the fact that LGG shows higher survival than GBM, so genes that are more frequently mutated in LGG or GBM showing some degree of relation with survival would likely be significant in GBMLGG. Nevertheless, we noted that significant genes in GBMLGG are not necessarily significant or marginally significant in LGG or GBM (Supplementary Figure 18). Thus, the significance in GBMLGG seems to be a reflect in gain of power given the aggregated number of samples.

Surprisingly many cancer types do not show significant gene mutations. It is possible that the small sample size could affect some cancer datasets. For example, in Papillary thyroid carcinoma (THCA-SA), we analyzed only 15 samples. Nevertheless, in other cancers like breast, bladder, and skin cancer (BRCA, BLCA, and SKCM), no significant genes were found associated with survival time at FDR=0.333 even though that these datasets include more than 900, 400, and 340 samples respectively. This indicates that more focused analyses in different strata are needed in these cancer types. Another example is CLLE-ES where no genes significant were found using 218 patients, but some of the top ones (EGR2, ASXL1, NOTCH1, POT1, and NXF1) have been reported recently ^31^ using more than 450 patients. So it is likely that we are detecting fewer genes.

As a proof of concept, we focused in coding mutations, however, it is known that copy number alteration (CNA) are also related to survival ^29,31^. Thus, further analyses should focus also on CNA data.

Some mutational biomarkers have been proposed for few cancer types ^31^. In this context, we find that some genes previously identified using the ALRT may change its significance, that others may climb up in rank, that a considerable proportion of genes provides low-risk odds, and that hypermutated samples may influence the identification. These results suggest that novel or refined cancer biomarkers can be identified.

Based in our simulations and analysis in cancer data, we demonstrated that VALORATE is fast, precise, and accurate to estimate the p-value of the difference of two survival curves using the log-rank statistic even in cases when the number of subjects in survival groups are highly unbalanced. We conclude that VALORATE is a novel and useful tool in cancer genomics and other statistical analyses.

## Methods

### The VALORATE algorithm

It is assumed that there are two groups of individuals and that for each patient we know their follow-up time and whether that time represent an event (e. g. death, metastasis, recurrence) or not (censored). *n* represents the total number of individuals and *n*_1_ the individuals in the mutated group. There would be then *r* distinct ordered times and *j=1..r* represents each of these times. Let *R*_*j*_ be individuals at risk that have not yet presented the event and *R*_*1j*_ those at risk for the mutated group. In each time *j*, there would be *O*_*j*_ events (zero or more), and *O*_*1j*_ events for the mutated group. Under the null hypothesis of no difference between groups, *O*_*1j*_ is hypergeometric, so the expected number of events in the mutated group is *E*_*1j*_=R_*1j*_\**O*_*j*_/*R*_*j*_ ^25^. The log-rank statistic is then the sum of differences between the expected and the observed number of events ^25^ as

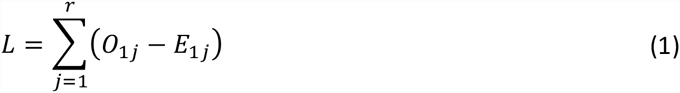

Under certain assumptions (high number of samples, not so few events, and similar group sizes ^26^), the mean of *L* is zero, and its variance is

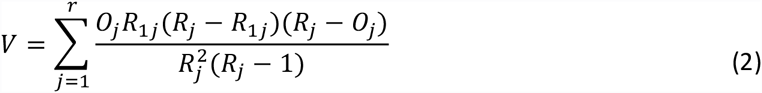

Thus, a *L^2^/V* follows a χ^2^ distribution with one degree of freedom and this fact can be used to estimate the p-value of *L* being zero equivalent to no difference in the survival curves. Nevertheless, the χ^2^ approximation yield bias estimations when *n*_1_ << *n/2* ^27^, which is the case in cancer genomics. To get accurate estimations of the probability of the two survival curves been equal when *n*_1_ << *n/2*, we first rewrite the equation (1) in terms of events ranked by time and their corresponding mutated group ^27^ as

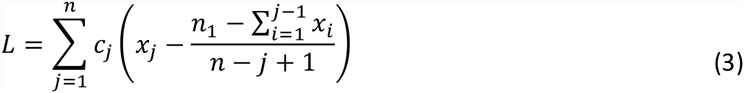

where *c*_*j*_ =*1* is the indicator of event (death, recurrence, metastases) or *c*_*j*_=0 for censored observations (events not yet observed) ordered by time, *x*_*j*_=1 for subjects that are included in the mutated group or *x*_*j*_ =0 for those who are not mutated, *n* is the total number of subjects, and *n*_1_, which is equal to *∑x_j_*, is the total number of subjects mutated. To estimate the permuted density of the test statistic *L*, we rearranged equation (3) as

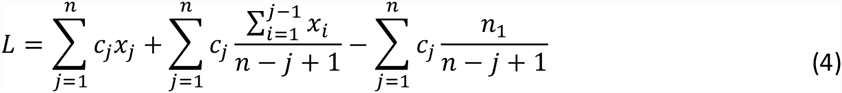

where it is evident that when *n*_1_≪ *n*/2, the Log-Rank statistic *L*, should depend strongly on the left term, 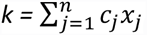, the number of co-occurrences, which represent events (*c*_*j*_ =1) that are also mutated (*x*_*j*_=1). The middle term also depends on *x_j_* but it is more robust to precise positions of *x*_*j*_=1 than the left term, which is highly dependent on the positions of *x*_*j*_=1. The right term, let be *s*_1_, is constant. From (4) it follows that *k-s_1_< L < k* for a particular value of *k*. This observation is important because it points out that the overall log-rank distribution can be seen as a mixture of distributions that depend on the number of co-occurrences *k*. The number of combinations for each co-occurrence *k*, although highly variable, can be easily calculated. To estimate a p-value, the relative proportion of combinations of the co-occurrences *k* is used to weight the relative contribution of the middle term to the overall distribution. Furthermore, the distribution of *L* conditional to a specific value of *k* (*L*_*k*_ = P(L|k)) can be estimated by random sampling (permutations) instead of an non-conditioned all-combinations approach used by other methods ^32^. Consequently, the VALORATE algorithm (Figure 1) estimates the distribution of *L* by a weighted sum of conditional distributions *L*_*k*_ for specific values of *k* such as *L= ∑w_k_L_k_*. Where *k* varies from 0 to *min(n_1_, d)* and *w*_k_ is the proportional contribution of the combinations for particular co-occurrences *k*. Finally, the probability of a specific observed statistic, *L_(gene)_*, is estimated by the area of the right (or left) fraction. The area is easily estimated summing the number of random samples of each *L*_*k*_ that are greater (or lesser) than *L*_*(gene)*_ multiplied by its corresponding weight *w*_*k*_. Ties can be broken by random sampling the ***c*** vector in tie positions during the estimation of *L_k_* only in tie positions containing mixtures of events and censored observations. During the estimation of the p-value, we estimate the average log-rank statistic permuting tie positions.

The main parameter for VALORATE is the total number of samples (*ss=sampling size*) used for the estimation of the whole distribution. For each value of *k*, the sampling size *ss*_*k*_ is obtained weighting *ss* respect to the probability of observing *k* (or a minimum of sampling, *ss*_*min*_, or all combinations if *ss_k_* is more than half of the number of combinations for *k*). We commonly use *ss=100,000*, and *ss_min_=1,000*. For each cancer dataset, this procedure was used to obtain *L* for each observed value of *n_1_* (usually for *n_1_ > 3* or *n_1_=3* in special cases, see the next sections). The p-values have to be multiplied by a factor of 2 for two-tailed tests, which was used for comparisons with the ALRT.

### Cancer mutation data

The mutation and clinical data were obtained from data portals. From TCGA (https://tcga-data.nci.nih.gov) specifically from the FireBrowse interface (http://firebrowse.org/) and from ICGC (https://dcc.icgc.org). A summary of the used data is shown in Supplementary Table 1. Overall, 61 cancer datasets were analyzed (35 from TCGA and 26 from ICGC) covering 12,428 cancer samples and 2,772,613 gene mutations (a gene may be mutated more than one time per sample). Only mutations carrying a clear coding effect (missense, nonsense, frame-shift, insertions, deletions, and splicing changes) were used avoiding mutations in introns and untranslated regions.

### Simulations of survival data and mutations

To determine the accuracy and precision of VALORATE, we performed simulations generating random ***c*** vectors for specific values of *n*, *n_1_*, and *d*. All possible values of ***L*** were calculated corresponding to all possible combinations of the ***x*** vector. Thus, the *exact* distribution of *L* was obtained and compared to the distribution estimated by VALORATE.

### Performance analyses

To evaluate and compare the running time of VALORATE, we used the glioblastoma dataset reported in the supplementary data from Vandin *et al.* ^27^. Specifically, we used the file tableR.txt obtained from https://github.com/fvandin/ExaLT. The p-value estimations used for comparisons with VALORATE in Figure 2 used all genes. From the 3 p-values (LEFT, RIGHT, SUM) provided by ExaLT, we used the closest compared to VALORATE. The estimations of CPU-time were performed for 4 genes only having the largest number of mutations and that are estimated by the ExaLT algorithm (marked as FPTAS) using the default parameters. Genes used were PIK3R1, IDH1, ERBB2, and SYNE1 having 12, 10, 9, and 7 mutations respectively.

### VALORATE analyses

For the simulations, the *ss* parameter used is specified in each particular experiment and *ss*_*min*_ was *1,000* unless specified. For the cancer data analyses, we used *ss=100,000*. We focused on genes mutated in more than 4% of the patients. Thus, for 46 cancer datasets having 75 or more subjects with survival data, only genes mutated in 4 or more subjects were used. For the 15 cancer datasets having 74 or fewer subjects, only genes mutated in 3 or more subjects were used.

### Selection of hypermutated samples

The range of mutation rates per cancer type is dependent on particularities of each cancer type ^10^. For instance, the median of the number of mutated genes of neuroblastoma (NBL-US) and thymus cancer (THYM) are 1 and 9 respectively, while this number in bladder (BLCA) and melanoma (MELA-AU) is 169 and 343 (Supplementary Figure 12). Accordingly, to avoid removing samples in cancer types having few mutations, hypermutated samples were removed if they have more than 500 mutated genes, are within the top 5% of most mutated samples, and the number of mutated genes is larger than the median plus four times the median absolute deviation. The specific number of samples removed per cancer type is shown in Supplementary Figure 12.

### Functional analyses of mutated genes

To validate whether the significant genes obtained by VALORATE could have a functional effect, we performed two analyses. First, we reasoned that a significant gene will likely be expressed to exert a functional role. Thus, we obtained the gene expression levels from TCGA or ICGC data portals of available cohorts to assess the overall relative expression of the significant genes including the comparison between all subjects and those mutated. Microarray or RNA-Seq data was used (Supplementary Table 1). Second, we thought that significant mutations should be more ‘damaging’ than random mutations. Therefore, we used MutationAssessor ^15^ to qualify the level of the functional impact of mutations. This tool classifies mutations according to evolutionary conservation patterns of affected amino acids in homolog proteins.

### Code

The VALORATE code was mainly implemented in R with some portions in C. VALORATE is freely available for download and usage under a general MIT license. Instructions for usage, example, and additional information can be found at https://github.com/vtrevino/valorate.

## Acknowledgments

We thank Arturo Berrones and Edgar Vallejo for discussions and comments.

## Author contributions

J.T.P. and E.M.L. contributed to method design, revision and discussion of the results, and writing the manuscript. V.T. conceptualized the study, designed and implemented the method, performed the analyses, and wrote the manuscript.

## Funding

We thank CONACyT for grant 233489 FONSEC SSA/IMSS/ISSSTE and Tecnológico de Monterrey for funding the Grupo de Investigación con Enfoque Estratégico en Bioinformática y Dispositivos Médicos.

## Competing financial interests

The authors declare no competing financial interests.

## Supplementary Files

Supplementary Figures.pdf – Contains all supplementary figures and legends.

Supplementary Table 1 – Datasets.xlsx – Contains the details of cancer datasets used.

Supplementary Table 2 – Gene Estimations.xlsx – Contains the p-value estimations of mutated genes across datasets.

## Supplementary Figure Legends

**Supplementary Figure 1. Comparisons of exact and estimated log-rank distribution for varied simulations.** The black line represents the exact distribution whereas the colored lines show the distributions estimated by VALORATE. The values of *n, d (ev), n1,* and a total number of combinations (*comb)* is included in the top of each panel. The top 8 panels correspond to the distribution of 8 simulated scenarios shown in nominal units. The bottom 8 panels display corresponding distributions in logarithm base 10 scale to highlight local modes of low density.

**Supplementary Figure 2. QQ plot comparison of distributions.** The distributions correspond to the simulation shown in Figure 2 of the main paper. The exhaustive distribution is shown in horizontal axis while the VALORATE distribution is shown in the vertical axis. Each dot corresponds to the value of the distribution from 0% to 100% in increments of 1%. The extreme dots around +/− 6 marked with arrows were seen in the exhaustive calculation but not observed in the random sampling of VALORATE, which is expected due to random nature of the sub-sampling process.

**Supplementary Figure 3. Comparisons of p-value estimations.** Each row of panels shows a specific simulation varying *n_1_={50, 30, 14, 7}* respectively randomizing the mutational group (*x* vector) 50,000 times and using *n=100* subjects and *d=10* events. The left column shows the observed distribution of *k* co-occurrences (death and mutations), followed by the p-value estimations in linear and logarithmic scales, and the overall *L* distribution estimated by VALORATE. The ALRT p-values are shown in the horizontal axis whereas the VALORATE p-values are shown in the vertical axis. Note that p-value differences are dependent on *n_1_* and *k*. Some co-occurrences were missing within the 50,000 random vectors in n_1_=14 and n_1_=7.

**Supplementary Figure 4. Examples of differences in the p-value estimation.** (A) Shows the estimated p-values from the ALRT (horizontal axis) and VALORATE (vertical axis) for a simulation having *n=100, d=10,* and *n1=7* (as in Supplementary Figure 3). Colors correspond to the number of co-occurrences (black=0, red=1, green=2, blue=3, cyan=4, magenta=5, and 6 and 7 were not observed in this sampling). “*” at the bottom right (black) and top left (red) marks two extreme cases shown in (B) and (C) respectively. (B) The estimated p-value using VALORATE of 1.8×10^−4^ which was estimated by the ALRT as p=0.15. (C) The estimated p-value using VALORATE of 0.27 which was estimated by the ALRT as p=3.5×10^−6^

**Supplementary Figure 5. Precision of VALORATE at different values of sampling size.** Two parameters sets (scenarios) were used. The top 2 rows show simulations at *n=100, d=10 (ev), n_1_=7* and the 2 bottom rows at *n=300, d=30 (ev), n_1_=4*. Columns show different values of the sampling size parameter (*ss*) corresponding to 10^3^, 10^4^, 10^5^, and 10^6^. Each panel shows two runs in different colors. Row 1 and 3 correspond to raw scale whereas rows 2 and 4 correspond to logarithm base 10 scale to highlight low-density regions.

**Supplementary Figure 6. Differences of p-value estimations across cancer types.** Each panel shows a cancer type, the samples used, the number of censored samples, the average number of mutations per sample, and the p-value estimations for VALORATE (horizontal axis) and the ALRT (vertical axis). Each dot corresponds to a gene in the dataset. Only genes whose p-value < 0.01 in any test and having 4 or more mutations are colored.

**Supplementary Figure 7. Differences of p-value estimations along a number of mutations.** Each panel shows the p-value estimated in VALORATE (horizontal axis) and the ALRT (vertical axis) for the specified number of samples mutated (from 3 to more than 25). Colors correspond to cancer types.

**Supplementary Figure 8. Number of significant genes at FDR=0.333 across cancer types.** (A) Significant genes using all samples in VALORATE and the ALRT. (B) Significant genes after removal of top 5% most mutated samples.

**Supplementary Figure 9. Comparison of significant and top genes.** (A) q-value of genes significant at q-FDR < 0.333 and p < 0.05 in VALORATE (horizontal axis) or in the ALRT (vertical axis). (B) Ranks of genes in (A).

**Supplementary Figure 10. Association of the number of significant genes with the numbers of samples.** Association to (A) the number of samples used, (B) deaths, (C) censored, and (D) percentage of censoring.

**Supplementary Figure 11. Association of the number of significant genes with hypermutated of samples. The** The panels show the top 30 most significant genes (having lowest p-values) in the vertical axis and samples in the horizontal axis. The number of mutations per subject and the survival days is also shown on top of each panel. The ordering of columns corresponds to the significance and whether the gene was associated with low risk (green) or high risk (red). The columns were ordered by the number of mutations in low and high risk within the genes shown.

**Supplementary Figure 12. Number of hypermutated samples removed along with mutated genes per sample and cancer type.** Each dot corresponds to a sample within a cancer type (horizontal axis). Cancer types ordered by the median of the number of mutated genes. The number of samples of each cancer type is shown in parenthesis. The number of ‘hypermutated’ samples removed are shown above the line marking the cut-off used. For most cancer types, a 500 cut-off value was used.

**Supplementary Figure 13. Number of significant genes per cancer type after removal of hypermutated samples.**

**Supplementary Figure 14. Significant genes using VALORATE in gliomas are not related to most mutated samples.**

**Supplementary Figure 15. Risk group associated with significant genes using VALORATE.**

**Supplementary Figure 16. Top genes seem cancer-type specific.** A) Significant genes relaxing the FDR cut-off to FDR < 0.999, p < 0.05, and maximum 10 genes per cancer type. (B) The p-value of the top 5 genes per cancer type.

**Supplementary Figure 17. Functional impact of the mutations in significant genes** (A) Functional impact of the mutations in significant genes. (B) Functional impact of random genes having a similar number of mutations within the same cancer types. The annotations were obtained from MutationAssessor. (C) Statistical analysis of the differences in functional impact.

**Supplementary Figure 18. Comparison of the significance of genes between Gliomas and Glioblastoma and Low-Grade Gliomas.** The figure shows the p-value estimated by VALORATE in logarithm base 10 scale for Glioblastoma (GBM) in the vertical axis, for Low-Grade Gliomas (LGG) in the horizontal axis and for Gliomas (GBMLGG) in the size of the bubble. Top genes in gliomas (GBMLGG) are labelled.

